# Quantification of spliced and unspliced transcripts by velocyto is inaccurate for 5’-sequencing data

**DOI:** 10.1101/2025.01.17.633503

**Authors:** Doreen Klingler, Jan T. Schleicher, Manfred Claassen

## Abstract

RNA velocity allows for the prediction of future states of individual cells based on their current transcriptional activity. The technique depends on the separate quantification of spliced and unspliced transcripts in single-cell RNA-sequencing data, first introduced by velocyto in 2018. Since its introduction, significant advances have been made in the field, including new protocols by 10x Genomics that enable sequencing from the 5’-end of mRNA molecules instead of the 3’-end. Despite these advances, velocyto has not been updated since its release but is still commonly used with these new protocols. In this study, we demonstrate that velocyto cannot accurately detect the reversed direction of transcripts, leading to incorrect count assignments. By comparing velocyto to alevin-fry, a quantification method compatible with 5’-sequencing data, we show that this limitation can result in substantial deviations in inferred velocities and differing interpretations. Therefore, we do not recommend the use of velocyto with 5’-sequencing data.

## 1 Introduction

Single-cell RNA sequencing (scRNA-seq) has enabled the study of dynamic processes of individual cells such as differentiation, cell cycle, and immune response [Saliba et al., 2014, Griffiths et al., 2018, Kulkarni et al., 2019]. The widely applied scRNA-seq platform 10x Genomics Chromium provides a range of protocols to sequence thousands of single cells simultaneously [Zheng et al., 2017]. Depending on the purpose of the scRNA-seq experiment, sequencing can capture either the 3’– or the 5’-end of the transcripts. Both approaches include the separation of individual cells into droplets, unique tagging of each mRNA molecule by a cell barcode and a unique molecular identifier (UMI) during reverse transcription, followed by library preparation and amplification. The final sequencing library consists of a read including the cell barcode and UMI (R1), and a second read containing the captured mRNA sequence (R2), ready for Illumina paired-end sequencing.

Multiple methods have been developed for the subsequent alignment of sequencing reads to a reference genome or transcriptome, including 10x Genomics’ Cell Ranger based on STAR [Dobin et al., 2013] aligner, as well as multiple pseudoaligners requiring reduced computational resources like salmon [Patro et al., 2017] or kallisto [Melsted et al., 2021]. The resulting matrix of gene read counts per cell forms the basis for a variety of analyses. Although usually only a static snapshot of the transcriptome is captured, reads can be classified based on their alignment to exonic and intronic regions of transcripts into spliced, unspliced, or ambiguously assigned reads [La Manno et al., 2018]. Separate quantification of these reads, as introduced by velocyto [La Manno et al., 2018], results in distinct count matrices of spliced and unspliced molecules. While velocyto operates on reads that are already aligned, e.g., by Cell Ranger, the package salmon/alevin-fry [He et al., 2022] provides a workflow starting from raw sequencing reads and producing three count matrices with transcripts annotated as spliced, unspliced, or ambiguous. This information enables the estimation of so-called RNA velocity, i.e. the production rate of spliced mRNA molecules for a specific gene [La Manno et al., 2018]. By selecting all genes with informative velocities (velocity genes) and aggregating their velocities for each cell, cell-to-cell transition probabilities can be calculated, allowing to predict the future state of each cell.

Despite the considerable progress in scRNA sequencing methods since its release, velocyto remains one of the most frequently used tools to estimate spliced and unspliced counts for RNA velocity analysis [Gorin et al., 2022]. Consequently, a considerable number of studies also apply velocyto to 5’-sequencing data [Stewart et al., 2021, Mathew et al., 2021, Fu et al., 2023, McClory et al., 2023, Liu et al., 2022, Zhong et al., 2023, Argyriou et al., 2022], although the method was developed prior to the release of 5’-sequencing protocols and has not been updated since. Here, we aimed to assess whether velocyto can accurately assign reads from 5’ data to their correct originating strand, a critical requirement for the reliable quantification of spliced and unspliced transcript counts.

In this study, we re-analysed three publicly available scRNA-seq datasets, sequenced in the 5’-direction and originally processed by Cell Ranger and velocyto [Stewart et al., 2021, Mathew et al., 2021, Fu et al., 2023]. We show that velocyto significantly underestimates the total number of transcripts and incorrectly assigns gene counts to overlapping genes of the reverse strand by assuming the opposite direction of sequencing reads. Moreover, we demonstrate that the resulting velocity estimations considerably differ from those based on the alevin-fry workflow, which can differentiate between 3’– and 5’-sequencing.

## 2 Results

### 2.1 Spliced, unspliced, and total counts estimated by velocyto significantly deviate from alevin-fry and Cell Ranger

After aligning raw scRNA-seq reads to the respective transcriptome by Cell Ranger or salmon, two tools for the quantification of spliced and unspliced transcripts were investigated: velocyto and alevin-fry. We compared the sum of spliced, unspliced and ambiguous counts of each cell, obtained by velocyto and alevin-fry, to the total counts identified by Cell Ranger (Fig. 1A). While the distribution of total counts per cell estimated by alevin-fry resembled the distribution of Cell Ranger counts, velocyto yielded significantly lower counts per cell, especially for the R2-only aligned samples of all three datasets. This became even more apparent when looking at the distribution of per-cell differences in total counts obtained from velocyto or alevin-fry, compared to Cell Ranger (Fig. 1B). Similarly for the paired-end aligned samples, the count differences of velocyto were significantly higher compared to alevin-fry, albeit being lower than those seen for the paired-end alignment.

**Fig. 1.**
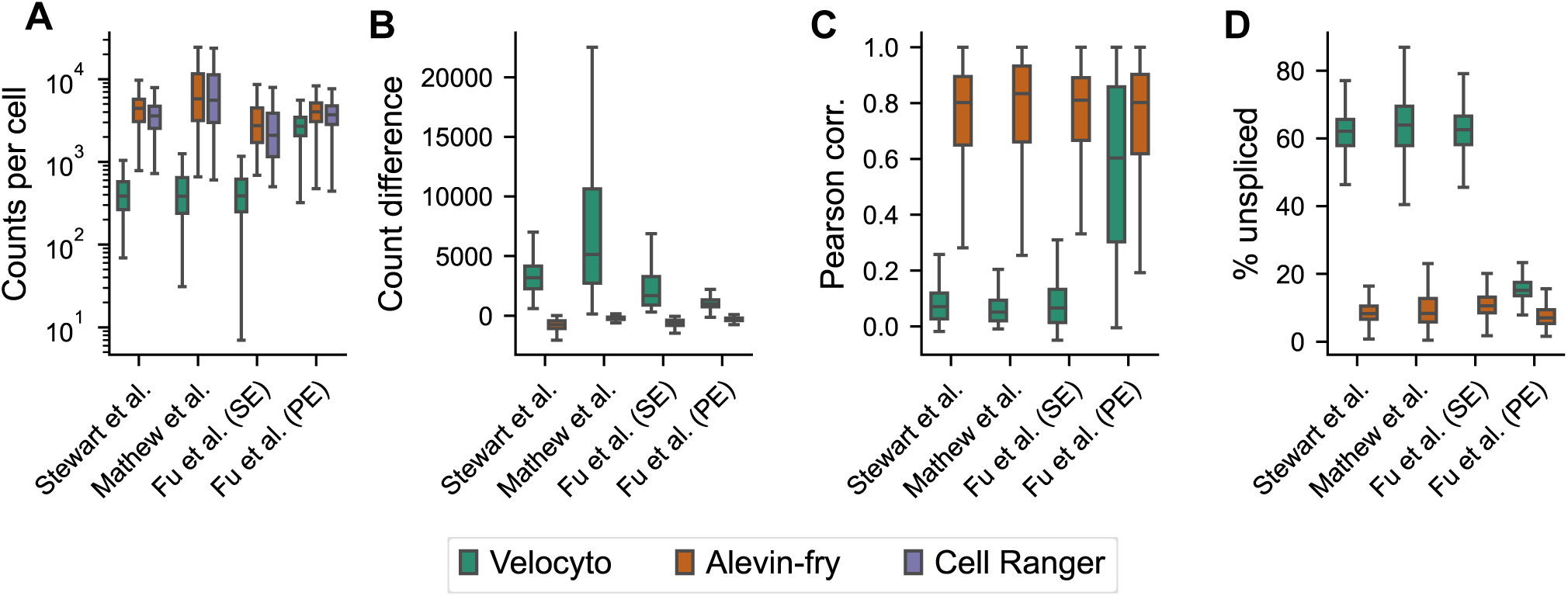
Comparison of various metrics based on the raw count matrices reveals strong disagreements between velocyto, alevin-fry and Cell Ranger. (A) Total gene counts per cell obtained from Cell Ranger and sum of spliced, unspliced and ambiguous counts for velocyto and alevin-fry, displayed for each dataset individually. The dataset from Fu *et al*. is additionally separated into samples aligned by single-end (SE) and paired-end (PE) alignment. (B) Difference of total counts per cell between velocyto/alevin-fry and Cell Ranger. (C) Pearson correlation of Cell Ranger counts with velocyto/alevin-fry spliced counts of the same gene. (D) Proportion of unspliced counts in the sum of spliced and unspliced counts per cell for velocyto and alevin-fry.

Furthermore, the mean Pearson correlation of Cell Ranger counts of a gene with its spliced counts was below 0.09 for all three datasets aligned by Cell Ranger using only R2 and quantified by velocyto (Fig. 1C). With paired-end alignment, the correlation was higher on average, with a mean of 0.57. Alevin-fry spliced counts showed a mean correlation of above 0.76 in each dataset.

We also compared the ratio of unspliced and spliced molecules per cell between velocyto and alevin-fry (Fig. 1D). While for alevin-fry, the mean percentage of unspliced transcripts was between 9 and 16% for all datasets, velocyto suggested much higher unspliced proportions with mean values above 64% for the R2-only aligned samples and around 20% for the paired-end aligned samples. La Manno et al. [2018] reported 15-25% of unspliced reads in scRNA-seq datasets.

### 2.2 Velocyto incorrectly assigns unspliced counts to overlapping genes

Next, we investigated the mechanistic causes of these discrepancies. Since velocyto does not take the orientation of reads into account, which differs between 5’ and 3’ protocols, we hypothesised that the observed unspliced counts do not arise from the genes themselves, but rather from transcripts on the opposing DNA strand at the same position. Indeed, we saw a high correlation of unspliced gene counts assigned by velocyto with the Cell Ranger counts of the respective overlapping gene (Fig. 2A, Fig. S1 and Fig. S2). The degree of correlation increases with greater overlap. (Fig. 2B). Moreover, the correlation did not originate from co-expression of the genes (Fig. S3), neither was it observed when estimating the counts by alevin-fry (Fig. S4). Correlations are also considerably lower when using Cell Ranger for paired-end alignment (Fig. 2B). For a small number of genes, counts were assigned to the spliced transcript of the overlapping gene (Fig. S5A), occurring when the overlap was located in exonic regions (Fig. S5B).

**Fig. 2.**
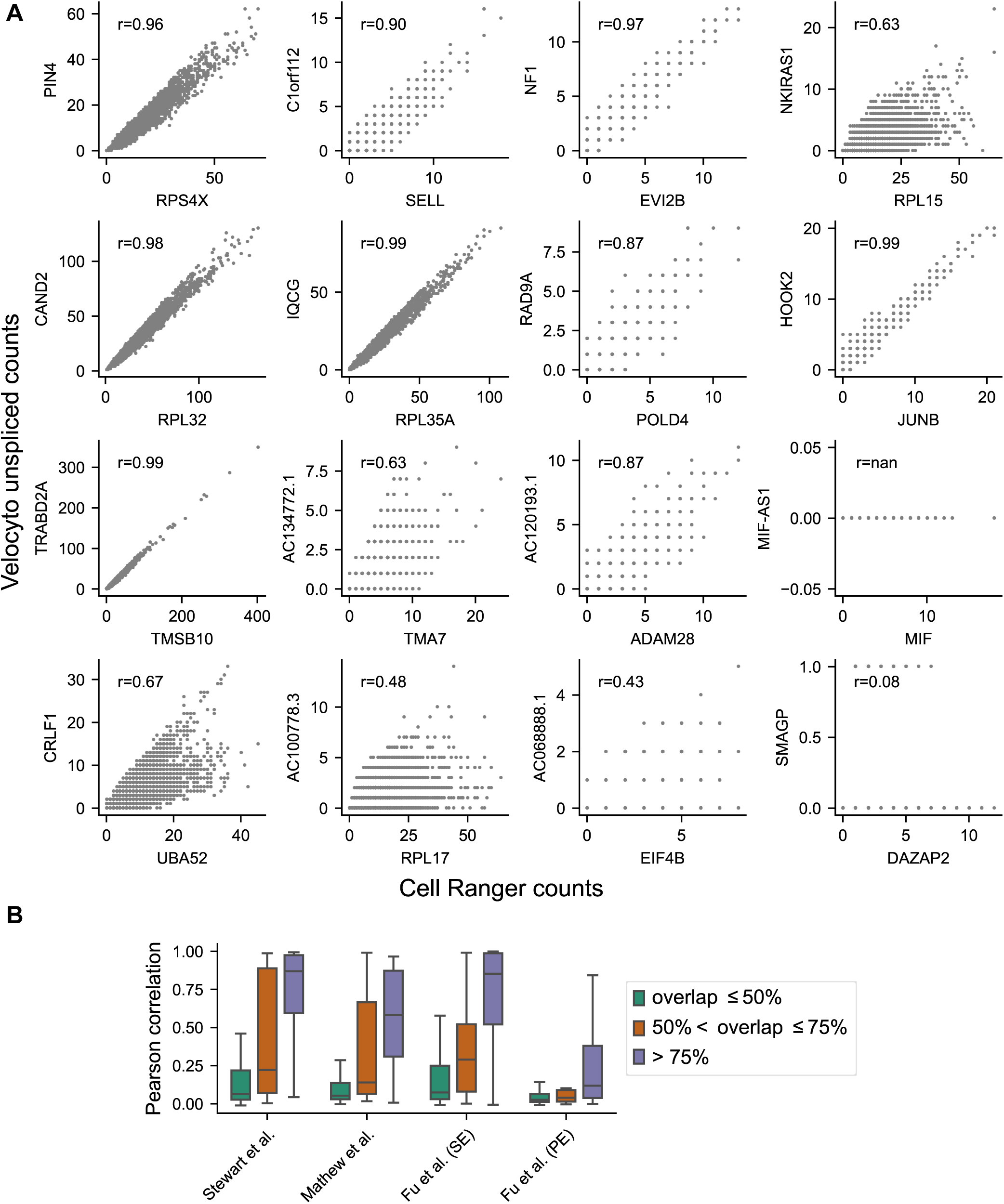
Velocyto incorrectly assigns unspliced counts to overlapping genes. (A) Correlation of Cell Ranger counts and velocyto unspliced counts of overlapping genes on opposite strands, exemplified by the data from Stewart *et al*. (r = Pearson correlation). Genes on the x-axis are sorted in descending order based on the proportion of their nucleotides that overlap with a gene located on the antiparallel strand (displayed on the respective y-axis). (B) Pearson correlation of Cell Ranger counts and velocyto unspliced counts of overlapping genes on opposite strands. Genes are grouped by the proportion of their nucleotides that overlap with a gene located on the antiparallel strand. The dataset from Fu *et al*. is separated into samples aligned by single-end (SE) and paired-end (PE) alignment.

### 2.3 Parameter estimation of velocity model strongly differs between velocyto and alevin-fry

As both the total counts per cell and per gene, as well as the unspliced counts and their ratio compared to spliced counts, strongly differed between velocyto and alevin-fry, we next studied the impact of this on RNA velocity estimation. First, velocyto consistently reported fewer velocity genes than alevin-fry and thus used a smaller number of genes for the prediction of future cell states (Fig. 3A). This was due to fewer highly variable genes meeting the R-squared value threshold of 0.01, which is a condition to be selected as a velocity gene by scVelo (Table S1. Furthermore, using velocyto also led to the prediction of different velocity genes that were not selected when alevin-fry was applied instead.

**Fig. 3.**
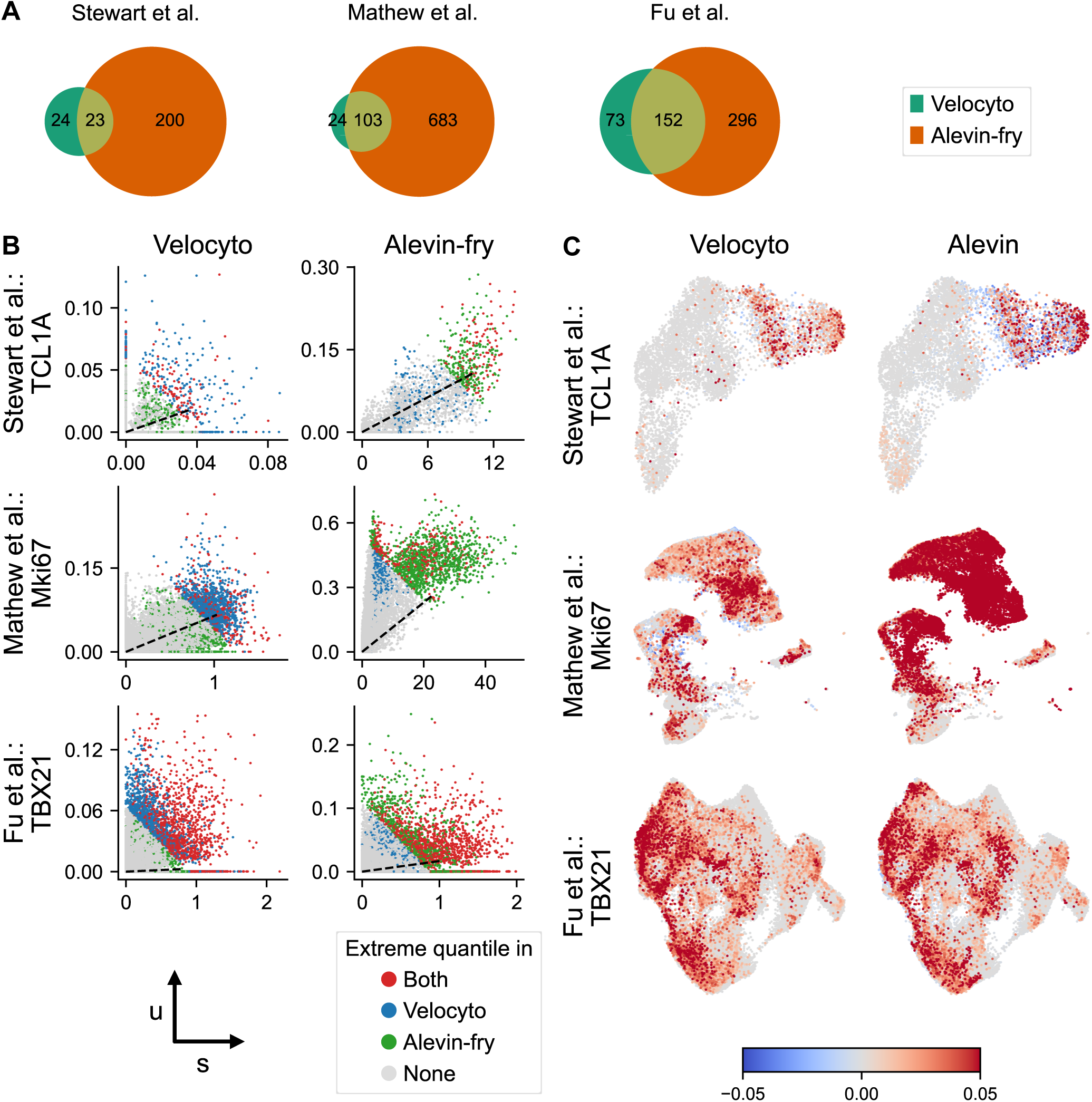
Comparison of velocity gene selection and velocity estimates. (A) Number and intersection of velocity genes based on velocyto and alevin-fry quantifications for the same datasets. (B) Phase portraits of marker genes with normalised spliced vs. unspliced counts, obtained from velocyto (left panels) and alevin-fry (right panels). Steady-state ratios (dashed black line) were calculated based on the stochastic model by a linear regression on extreme quantiles. Cells within the extreme quantiles are indicated by distinct colors, determined by the quantification method. For each dataset, one representative marker gene is shown, which was selected as velocity gene in both velocyto and alevin-fry quantifications. (C) Velocities of individual marker genes, visualised in the respective UMAP embedding. Velocities were estimated using the steady-state stochastic model and clipped to the interval between –0.05 and 0.05 for visualisation.

By comparing the phase portraits of important marker genes, we detected considerably lower normalised spliced counts in the datasets from Stewart et al. [2021] and Mathew et al. [2021] analysed by velocyto compared to alevin-fry (Fig. 3B). This discrepancy resulted in different cells being located in the extreme quantiles and therefore used for linear regression, which also caused deviations in steady-state ratios. This ultimately led to different positions of the same cells relative to the steady-state ratio, with the consequence that those cells were assigned velocities of different sign (Fig. 3C). While for velocyto, the majority of cells in the *TCL1A* phase portrait was located above the steady-state ratio and thus expected to be in the induction state, a more even distribution of cells in induction and repression state was observed for the alevin-fry data (Fig. 3B). Consequently, more cells with negative single-gene velocities were visible in the UMAP representation (Fig. 3C). The exact opposite applied to *Mki67*, again causing different velocity directions. These discrepancies were not observed for genes from Fu et al. [2023] data.

### 2.4 RNA velocities and their interpretation vary among the quantification methods

Next, we investigated and quantified how RNA velocities differed between velocyto and alevin-fry, and how this impacted their interpretation. We observed a mean cosine similarity of velocity vectors per cell below 0.1 for all datasets (Fig. 4A), indicating that velocity vectors for the same cells were almost orthogonal. Moreover, these disparities of velocity vectors were not related to a small number of discordant genes. In the R2-only aligned samples of Stewart *et al*. and Mathew *et al*., the majority of velocity vectors included more than 45% of genes with opposing velocities, while the mean proportion in the Fu *et al*. dataset was 25% (Fig. S6A). The substantial differences in velocity vectors between velocyto and alevin-fry were further validated by high cosine similarities between the velocity vectors of each cell and those of its nearest neighboring cell (Fig. S6B), with values exceeding 0.9 in all datasets. Additionally, the mean Pearson correlation of velocities per gene between velocyto and alevin-fry was below 0.25 for all datasets (Fig. 4B). Consequently, the transition probability distributions, forming the basis for the embedding of velocity vectors, also considerably differed between the two quantification methods, as indicated by a mean Wasserstein distance of more than 900 (Fig. 4C). Moreover, this divergence in transition probabilities is not only based on the selection of different velocity genes, but even using identical velocity genes could decrease the Wasserstein distance only slightly (Fig. S6C). Ultimately, discrepancies between velocities obtained from velocyto and alevin-fry, indicated by low cosine similarities and high Wasserstein distances are independent of clusters (Fig. S7A, C), as well as low Pearson correlations were observed both in highly variable and non-highly variable genes (Fig. S7B).

**Fig. 4.**
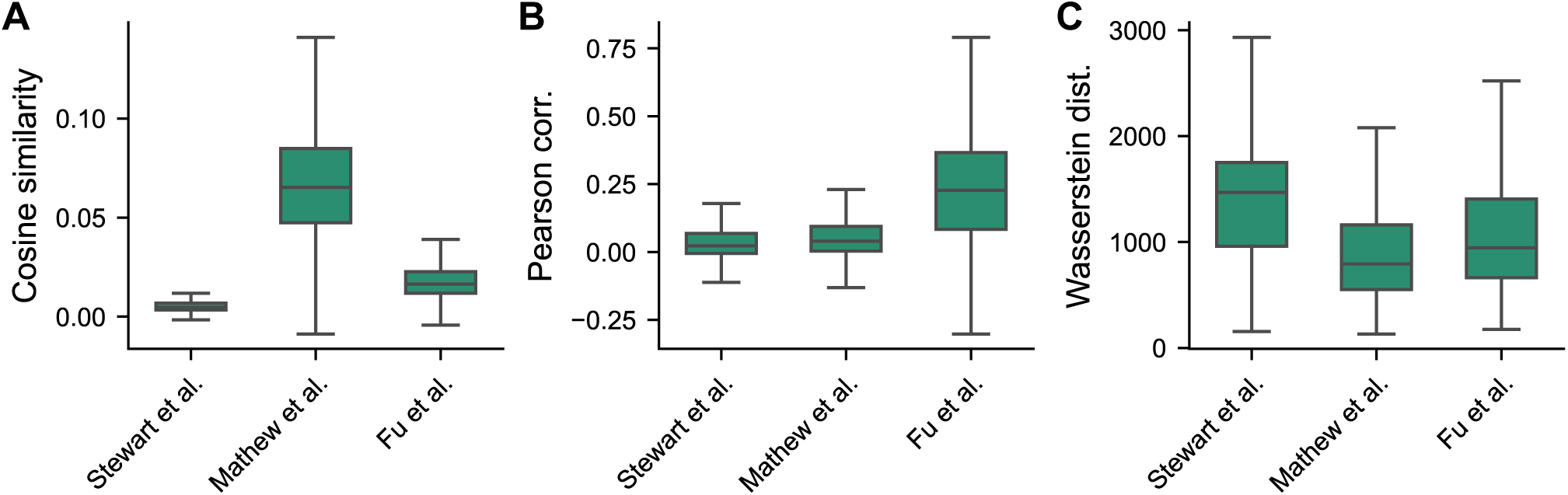
Comparison of velocities based on counts obtained from velocyto vs. alevin-fry. (A) Cosine similarity of velocity vectors per cell. (B) Pearson correlation of velocities per gene. (C) Wasserstein distance of transition probability distributions per cell.

Regarding the interpretation of RNA velocities, both in the 2-dimensional projection of velocity vectors, as well as in the visualisation of probable cell transitions by Partition-based graph abstraction (PAGA) [Wolf et al., 2019], deviating transition directions of individual cells or clusters were proposed when using velocyto, compared to alevin-fry (Fig. 5). For instance, in the dataset from Stewart *et al*., the projected velocity vectors distinctly point from the DN3 cluster to the C-mem2 cluster when using alevin-fry (Fig. 5B). In contrast, the vectors in the velocyto dataset predominantly point towards the DN4 and Naive clusters (Fig. 5A). Beyond that, more pronounced discrepancies are evident in the PAGA visualisation. While a transition from cluster DN1 to DN4 was predicted based on counts from velocyto (Fig. S5C), using alevin-fry resulted in the opposite direction (Fig. S5D). Similar discrepancies were also observed in other clusters and the other datasets (Fig. S8 and Fig. S9).

**Fig. 5.**
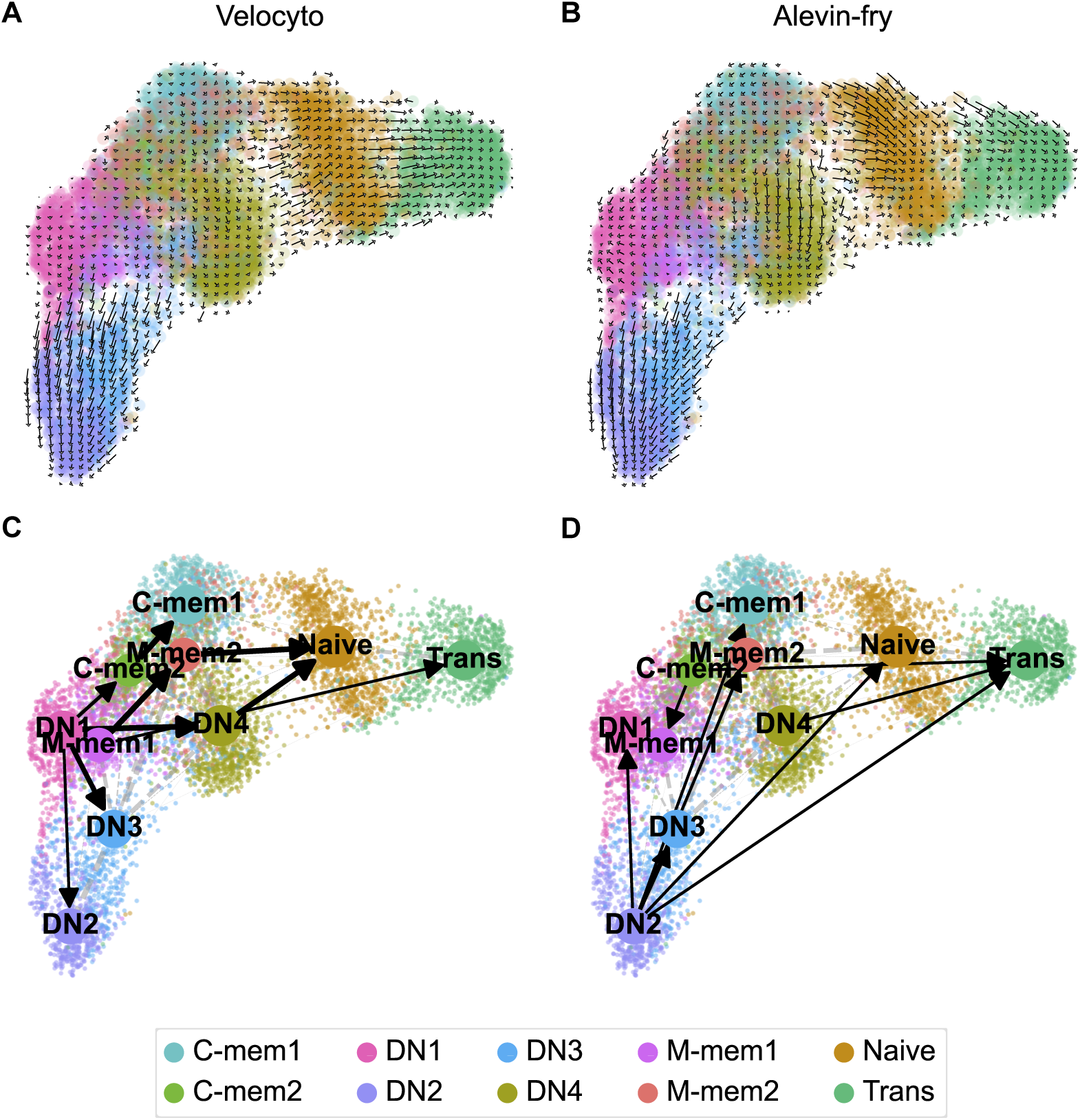
2-dimensional projections of velocities differ between velocyto and alevin-fry. (A, B) Visualisation of velocities from Stewart *et al*. data on a grid in the UMAP-embedding, with counts obtained from velocyto (A) vs. alevin-fry (B). (C, D) PAGA cluster relations projected on UMAP-embedding, with counts obtained from velocyto (C) vs. alevin-fry (D).

## 3 Discussion

Since velocyto has been developed using only 3’-sequencing data, we applied velocyto on three 5’-sequencing scRNA-seq datasets and compared the results to alevin-fry, which provides an option to specify the library orientation. We could demonstrate that velocyto wrongly assigns reads to overlapping genes on the opposite strand and overall reports considerably different counts for spliced and unspliced transcripts than alevin-fry, ultimately leading to deviating interpretations of RNA velocity.

Although differences in total counts between velocyto and alevin-fry have previously been reported in 3’-sequencing data, with a ratio of approximately 1.3 [Soneson et al., 2021], our analysis demonstrates that when applied to 5’-sequencing data aligned by Cell Ranger using R2-only alignment, velocyto reports on average 7 to 17 times fewer total counts compared to alevin-fry. Additionally, the majority of counts come from unspliced transcripts, which disagrees with the assumption of 15-20% of unspliced molecules by La Manno *et al*.. Paired-end alignment could reduce these discrepancies and the number of incorrectly assigned overlapping genes. However, for some genes, the assignment remained inaccurate, providing additional indications that velocyto should not be applied to 5’ scRNA-seq data at all.

Nevertheless, several studies performed RNA velocity analysis on 10x Genomics 5’-sequencing data, quantifying spliced and unspliced transcripts with velocyto [Stewart et al., 2021, Mathew et al., 2021, Fu et al., 2023, McClory et al., 2023, Liu et al., 2022, Zhong et al., 2023, Argyriou et al., 2022]. The number of such studies is increasing, for instance due to the compatibility of 5’ protocols with B/T cell receptor repertoire sequencing. This approach can have several critical implications, as evident from the low concordance in velocity estimates for identical cells or genes when analysed using velocyto and alevin-fry, as well as from the substantial discrepancies in transition probabilities. Accordingly, interpretations of cellular dynamics and biological processes become unreliable if velocities are inferred from inaccurate spliced and unspliced counts. Moreover, any subsequent analyses depending on RNA velocity, such as trajectory inference, pseudotime ordering, or the identification of regulatory networks, are likely to result in incorrect interpretations.

Despite these strong contradictions, the visualisations of velocity in 2-dimensional UMAPs exhibit only minor differences. This is potentially due to the fact that velocity vectors can only point to neighboring cells in this representation and are thus strongly influenced by the UMAP itself, as reported by Zheng *et al*. [Zheng et al., 2023].

Consequently, relying solely on two-dimensional visualisations for interpreting RNA velocity potentially poses notable risks, as it may lead to inaccurate or misleading conclusions.

In summary, we recommend against using velocyto for the quantification of spliced and unspliced reads from 5’-sequencing data and suggest an alternative workflow by alevin-fry. However, other quantification tools might be compatible with 5’-sequencing data, too, such as kallisto-bustools [Melsted et al., 2021] or dropEst [Petukhov et al., 2018], although they do not provide particular options to specify the sequencing directions and were therefore not included in this study. Moreover, the problem remains that there is no ground truth to evaluate RNA velocity methods. Therefore, a detailed understanding of the specific scRNA-seq protocols, the alignment algorithms, and the quantification methods is required to determine the best processing workflow and choose the right tools for each dataset individually. This also requires more studies revealing the bias of using different workflows and accounting for these differences when interpreting RNA velocities.

## 4 Materials and Methods

### 4.1 Data retrieval

Datasets used for this study are summarised in Table 1. For data from Stewart *et al*. and Mathew *et al*., FASTQ files of the raw sequencing reads were obtained from ArrayExpress by accession numbers E-MTAB-9544 [Stewart et al., 2021] and E-MTAB-9491 [Mathew et al., 2021]. The Fu *et al*. dataset was obtained by converting the BAM files deposited at NCBI SRA under accession number PRJNA922954 [Fu et al., 2023] back to FASTQ files using 10X Genomics’ bamtofastq. Moreover, to make results as comparable as possible to the original analysis, processed data from Stewart *et al*. and Fu *et al*. were obtained from the respective accession numbers and used as references. Code from Mathew *et al*. and Fu *et al*. was accessed from https://github.com/angelettilab/scMouseBcellFlu and https://github.com/princello/scRNA-seq-TRM-paper.

**Table 1.** Description of public datasets used for the study.

| Dataset | Organism | Cell type | # Samples | # Cells | # Genes | Alignment Method |
| --- | --- | --- | --- | --- | --- | --- |
| Stewart et al. [2021] | Human | B cells | 5 | 7,257 | 15,656 | R2-only |
| Mathew et al. [2021] | Mouse | B cells | 24 | 29,916 | 14,894 | R2-only |
| Fu et al. [2023] | Human | T cells | 12 | 50,033 | 20,408 | R2-only (5 samples)<br>Paired-end (7 samples) |

### 4.2 Alignment

Raw reads were aligned to the respective Ensembl reference transcriptome, human GRCh38 or mouse mm10, with Cell Ranger (v.7.1.0) command count. Assay chemistry was detected automatically, implying that the reads used for alignment depended on the sequencing length of R1. In seven out of twelve samples of the Fu *et al*. dataset, R1 was sequenced longer than 81 bases and thus both R1 and R2 were used by Cell Ranger for the alignment. For all other samples, alignment was performed only with R2, as the sequencing length of R1 was too short.

Splici references of the respective transcriptomes were generated by pyroe (v.0.9.3) according to the specific read length of each sequencing experiment (Stewart *et al*.: 100 bp, Mathew *et al*.: 110 bp, Fu *et al*.: samples MJ001, MJ002, MJ003, MJ016 and MJ017: 91 bp, samples MJ005, MJ006, MJ007, MJ008, MJ009, MJ018 and MJ019: 101 bp) with a flank trim length of 5 bp and used to build transcriptome index files with the salmon (v.1.10.2) function index. Next, raw reads were quantified against the respective index file by salmon alevin. This alignment approach always uses both R1 and R2.

### 4.3 Preprocessing

For each dataset, filtering and preprocessing were performed as described in the respective publication. All steps were applied identically on the Cell Ranger and salmon count matrices.

#### 4.3.1 Stewart *et al*

Count matrices were processed using scanpy (v.1.9.3). Cells were filtered to contain only B cells by selecting cells with zero counts of *CD3E*, *GNLY*, *CD14*, *FCER1A*, *GCGR3A*, *LYZ*, *PPBP* and *CD8A*. Furthermore, cells expressing less than 200 unique genes and cells with a total transcript count in the top 1% percentile were removed. Finally, only cells and genes present in both the filtered Cell Ranger and salmon data, as well as in the processed reference of Stewart *et al*., were selected. Gene counts were log-normalised and scaled with values clipped to a maximum of 10. Principal components (PC) were obtained from the reference. 14 PCs were used to calculate a neighboring graph of 20 nearest neighbors on the Cell Ranger dataset and the nearest neighbor graph was used for the salmon data as well. Stewart *et al*. performed Uniform Manifold Approximation and Projection (UMAP) on the first 14 PCs and used the Louvain [Blondel et al., 2008] algorithm to define clusters. For our analysis, UMAP positions and cluster assignments were obtained from the reference.

#### 4.3.2 Mathew *et al*

Data was processed following the Sauron (https://github.com/NBISweden/sauron) pipeline using Seurat (v3.0.1). This included removing all cells meeting one of the following criteria: percentage of mitochondrial or ribosomal gene counts >25%, expression of less than 200 distinct genes, number of unique features or total counts in the bottom or top 0.5% of all cells, or a Gini or Simpson diversity index <0.8. Additionally, mitochondrial genes, non-protein-coding genes, and genes expressed in fewer than 5 cells were filtered out, while immunoglobulin genes were kept. Next, gene counts were normalised, log-transformed, and scaled. The effects of total gene counts, percentage of mitochondrial counts, and G2M and S phase score difference per cell were regressed out. Per sample, 2,000 highly variable genes were selected and used to construct a neighboring graph and perform PCA. Samples were integrated based on the top 20 nearest neighbors using 51 PCs. Subsequently, the integrated dataset was reduced to 10 dimensions by UMAP, and Louvain [Blondel et al., 2008] clustering was performed based on these reduced dimensions. Cell types were predicted by analysing differential expression between clusters and the correlation of expression profiles with cell-type-specific gene signatures. Cells predicted to be non-B cells were removed and remaining cells were processed by the entire workflow again. Ultimately, 16 clusters were defined by hierarchical clustering based on 10 dimensions of the UMAP embedding. Clusters were annotated according to their expression of marker genes, as described by Mathew *et al*. [Mathew et al., 2021]. Finally, Cell Ranger and salmon datasets were subset to 29,916 common cells and 14,894 common genes. To minimise differences in the scRNA-seq data, highly variable genes, PCs, neighbors, cluster annotations, and UMAP coordinates calculated on the Cell Ranger data were transferred to the salmon dataset.

#### 4.3.3 Fu *et al*

scRNA-seq data was processed with Seurat (v.4.0.0), as described by Fu et al. [2023]. First, T cell receptor and immunoglobulin genes were discarded. Cells expressing >15% mitochondrial genes were removed, as well as cells with a unique gene count lower than the first quartile – 1.5 * interquartile range (IQR) or higher than the third quartile + 1.5 * IQR. Instead of random downsampling, which was performed in the original publication, all cells present in the reference were retained. Gene counts were normalised and scaled, and 6,400 highly variable features per sample were used for anchor-based data integration based on 64 PCs. The integrated datasets of Cell Ranger and salmon count data were further reduced to 50,033 cells and 20,415 genes present in the data of both workflows, as well as in the published data of Fu *et al*.. Moreover, 5,507 highly variable features, PCs, UMAP coordinates, and cluster assignments of 15 clusters, originally obtained by Louvain [Blondel et al., 2008] algorithm, were retrieved from the published data. A neighborhood graph of 20 nearest neighbors was constructed based on the Cell Ranger data with 30 PCs and also used for the salmon dataset.

### 4.4 RNA velocity

Spliced and unspliced transcripts were quantified by two methods. The BAM files of the Cell Ranger alignment were processed with veloyto (v.0.17.17) using velocyto run10x. The RAD files generated by salmon were further processed by alevin-fry (v.0.9.0). First, permit lists were generated by alevin-fry generate-permit-list and barcodes were corrected accordingly by alevin-fry collate. Ultimately, quantification was performed using alevin-fry quant. Spliced and unspliced counts and their ratio were compared between the two methods before normalising the counts. After count normalisation, RNA velocities were calculated by the steady-state model including second-order moments (stochastic model) using scVelo (v.0.3.2) [Bergen et al., 2020]. While the comparison of velocity vectors per cell and per gene between velocyto and alevin-fry data is not influenced by the selection of velocity genes, transition probabilities per cell are calculated using only velocity genes. Genes are selected as velocity genes if the following criteria are met: gene is highly variable, variance in the normalised unspliced counts explained by the linear regression model, determined by R-squared, >0.01, steady-state ratio >0.01 and normalised spliced and unspliced counts >0 in at least one cell. Transition probabilities reflect the cosine similarity of the cell’s velocity vector and the gene expression difference vector to the second cell. The resulting transition matrix was used to project velocities onto 2-dimensional UMAP embeddings on a grid. Additionally, predicted cluster transitions were calculated and visualised by PAGA [Wolf et al., 2019]. PAGA infers the connectivity between cells and clusters by integrating information from both cell clustering and pseudotemporal ordering. For pseudotime estimation, root cells were computed from the velocity-inferred transition matrices. For the dataset from Fu *et al*., PAGA was performed on a selected subset of groups (c01, c02, c03, c04, c07), as described in their analysis [Fu et al., 2023]. Gviz (v.1.48.0) was used to visualise gene overlaps [Hahne and Ivanek, 2016].

## 5 Data availability

All datasets used for this study are publicly available. Code with instructions to reproduce all analyses is available on GitHub.

## 6 Competing interests

No competing interest is declared.

## 7 Author contributions statement

Conceptualisation: J.T.S. and M.C., methodology: D.K. and J.T.S., analysis: D.K., writing: D.K., J.T.S., and M.C.

## 8 Acknowledgments

We thank the members of the Claassen group for constructive discussions and feedback. The authors thank the International Max Planck Research School for Intelligent Systems (IMPRS-IS) for supporting J.T.S. This work was supported by DFG CL 792/1-1.

## Supplementary Material

**Fig. S1.**
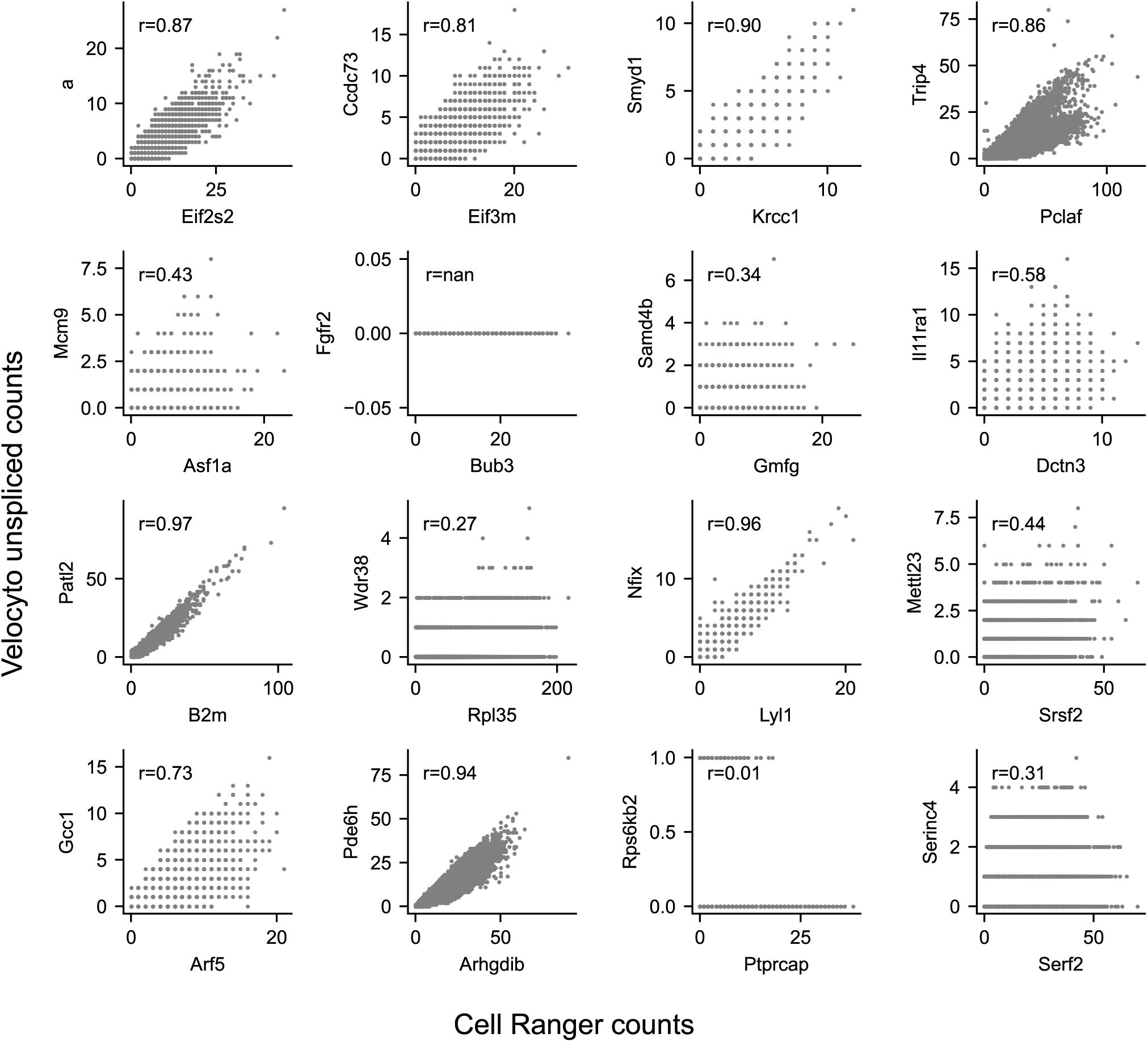
Correlation of Cell Ranger counts and velocyto unspliced counts of overlapping genes on opposite strands, shown for selected genes of the Mathew *et al*. dataset (r = Pearson correlation). Genes on the x-axis are sorted in descending order based on the proportion of their nucleotides that overlap with a gene located on the antiparallel strand (displayed on the respective y-axis).

**Fig. S2.**
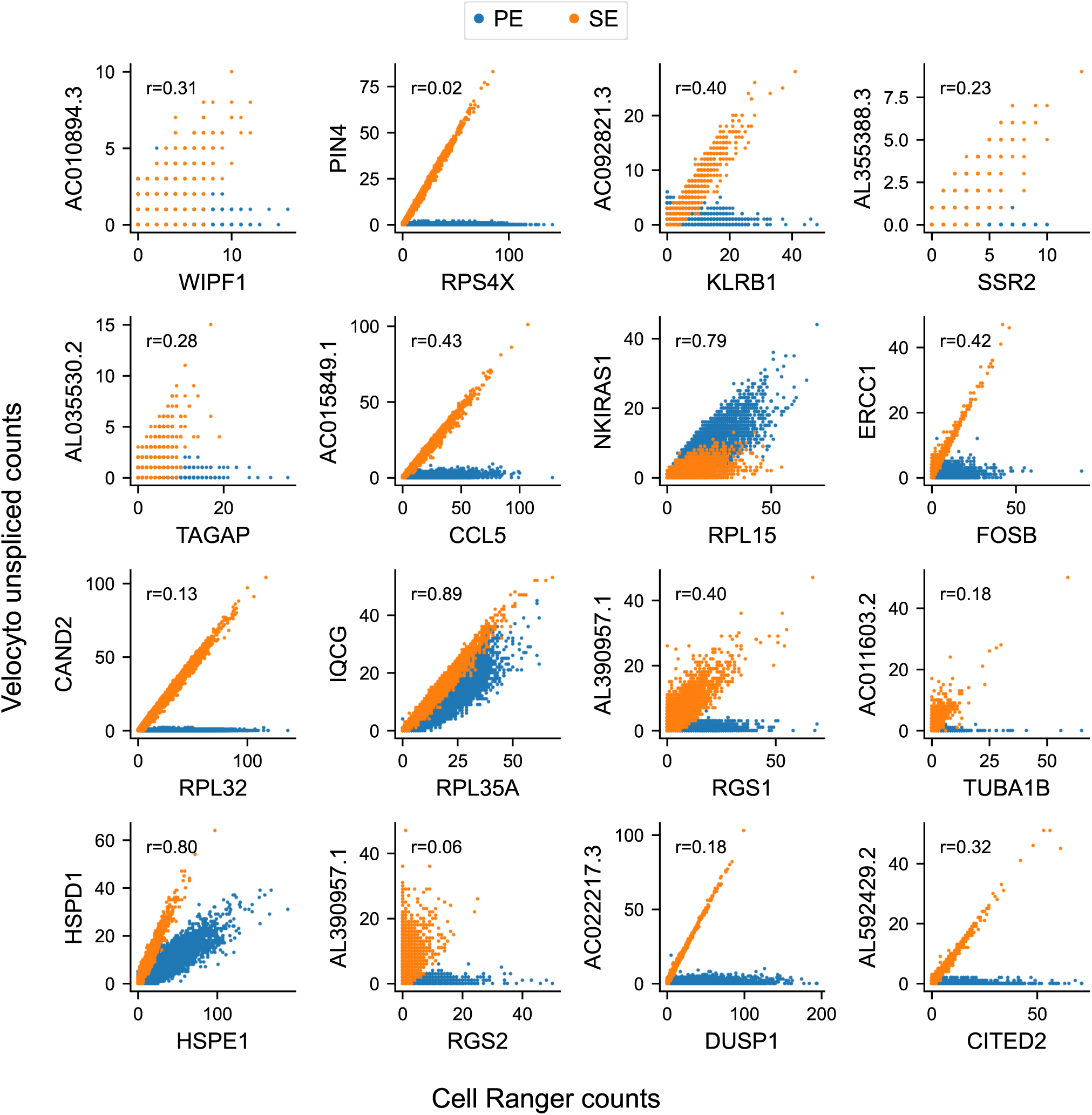
Correlation of Cell Ranger counts and velocyto unspliced counts of overlapping genes on opposite strands, shown for selected genes of the Fu *et al*. samples (r = Pearson correlation). Colors indicate Cell Ranger alignment mode (blue = paired-end alignment, orange = single-end alignment). Genes on the x-axis are sorted in descending order based on the proportion of their nucleotides that overlap with a gene located on the antiparallel strand (displayed on the respective y-axis).

**Fig. S3.**
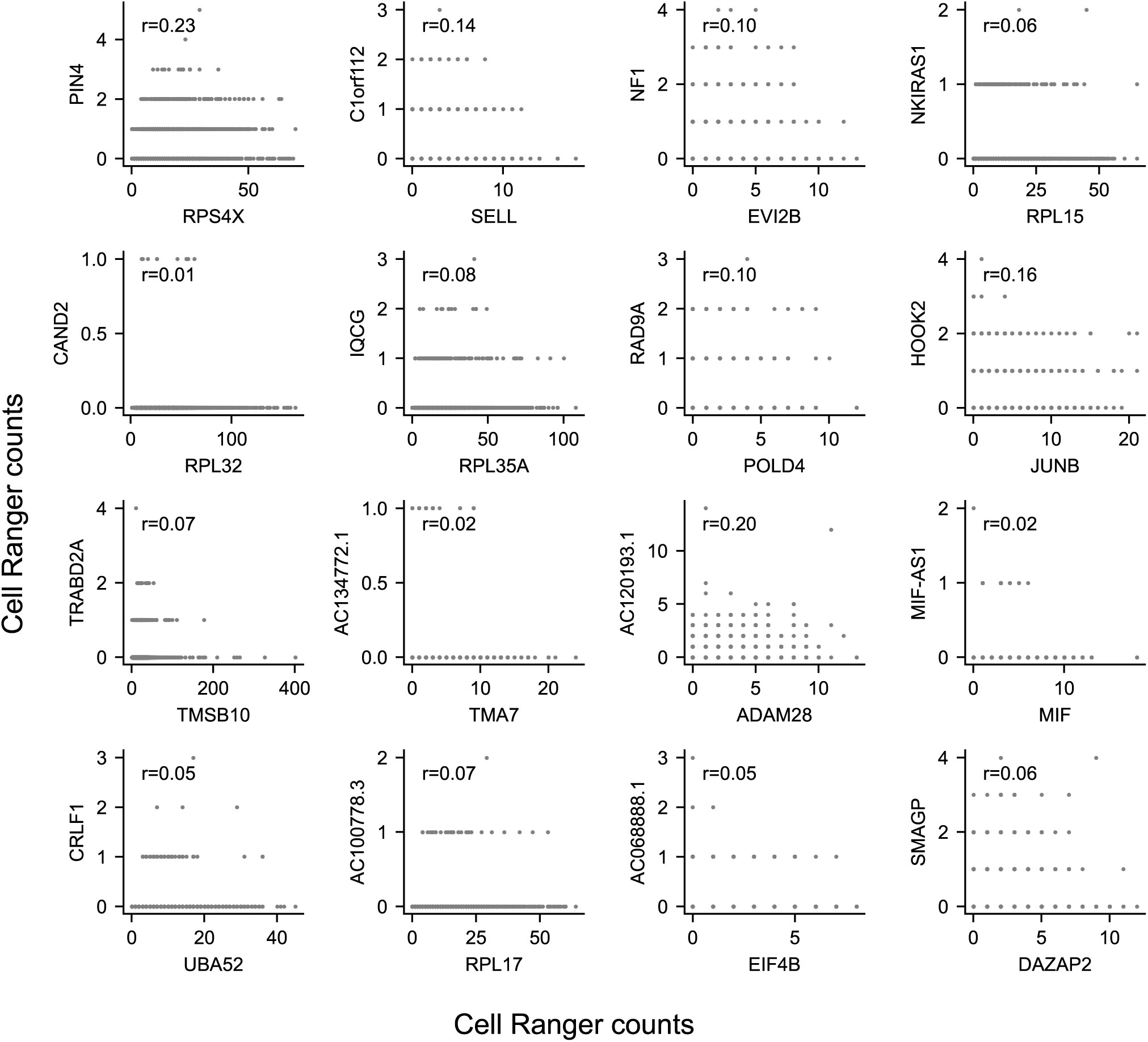
No co-expression observed between overlapping genes. Cell Ranger counts vs. Cell Ranger counts of overlapping genes on opposite strands, shown for the same genes of the Stewart *et al*. dataset as in Fig. 2. Genes on the x-axis are sorted in descending order based on the proportion of their nucleotides that overlap with a gene located on the antiparallel strand (displayed on the respective y-axis).

**Table S1.** Number of highly variable genes meeting the R-squared threshold of 0.01 to be selected as velocity gene by scVelo.

|  | Stewart *et al.* | Mathew *et al.* | Fu *et al.* |
| --- | --- | --- | --- |
| Velocyto | 47 | 128 | 248 |
| Alevin-fry | 240 | 802 | 571 |

**Fig. S4.**
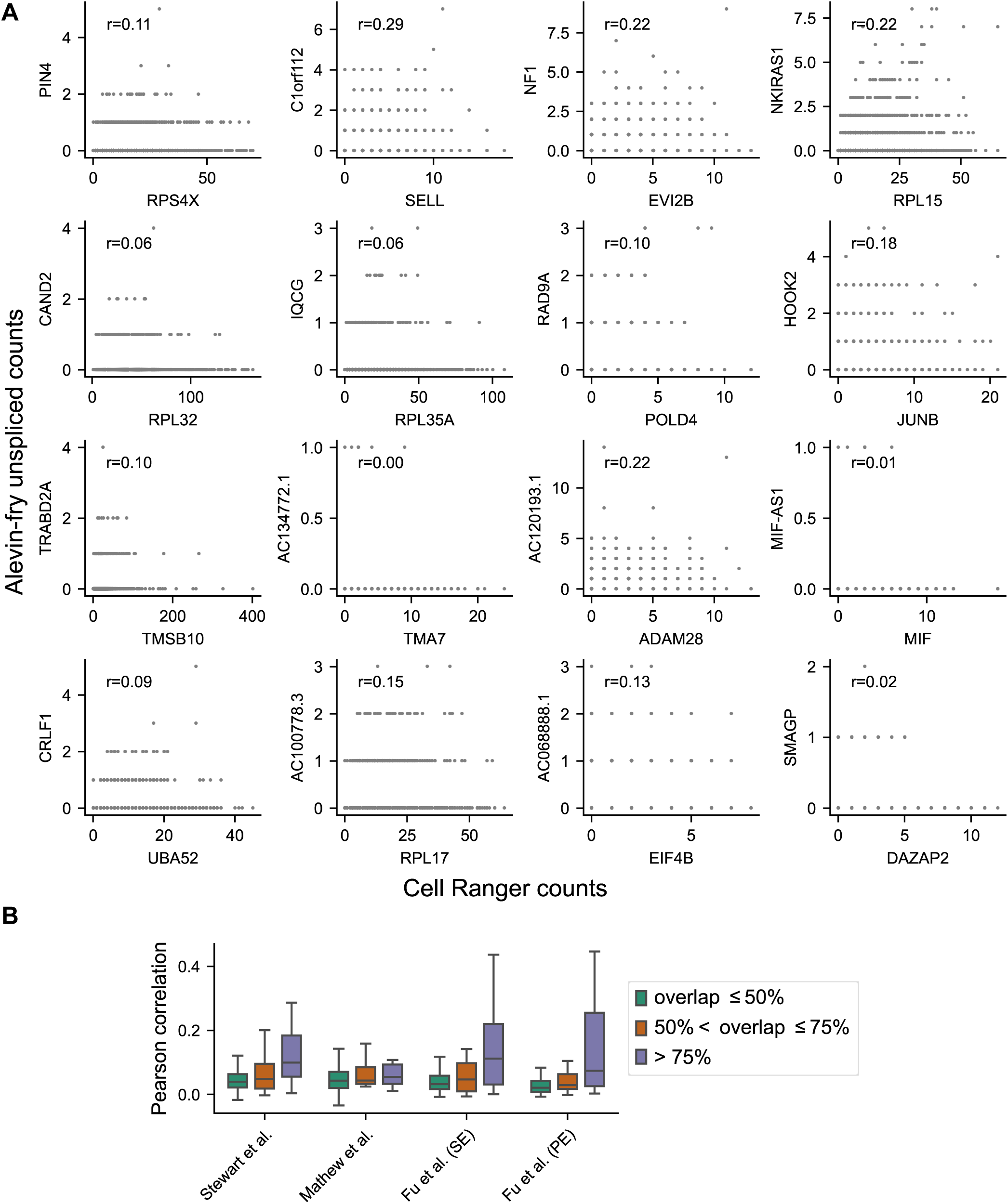
Alevin-fry correctly identifies direction of overlapping genes. (A) Correlation of Cell Ranger counts and alevin-fry unspliced counts of overlapping genes on opposite strands, exemplified by the data from Stewart *et al*. (r = Pearson correlation). Genes on the x-axis are sorted in descending order based on the proportion of their nucleotides that overlap with a gene located on the antiparallel strand (displayed on the respective y-axis). Similar patterns were observed in the data from Mathew *et al*. and Fu *et al*. with R2-only alignment. (B) Pearson correlation of Cell Ranger counts and alevin-fry unspliced counts of overlapping genes on opposite strands. Genes are grouped by the proportion of their nucleotides that overlap with a gene located on the antiparallel strand. The dataset from Fu *et al*. is separated into samples aligned by single-end (SE) and paired-end (PE) alignment.

**Fig. S5.**
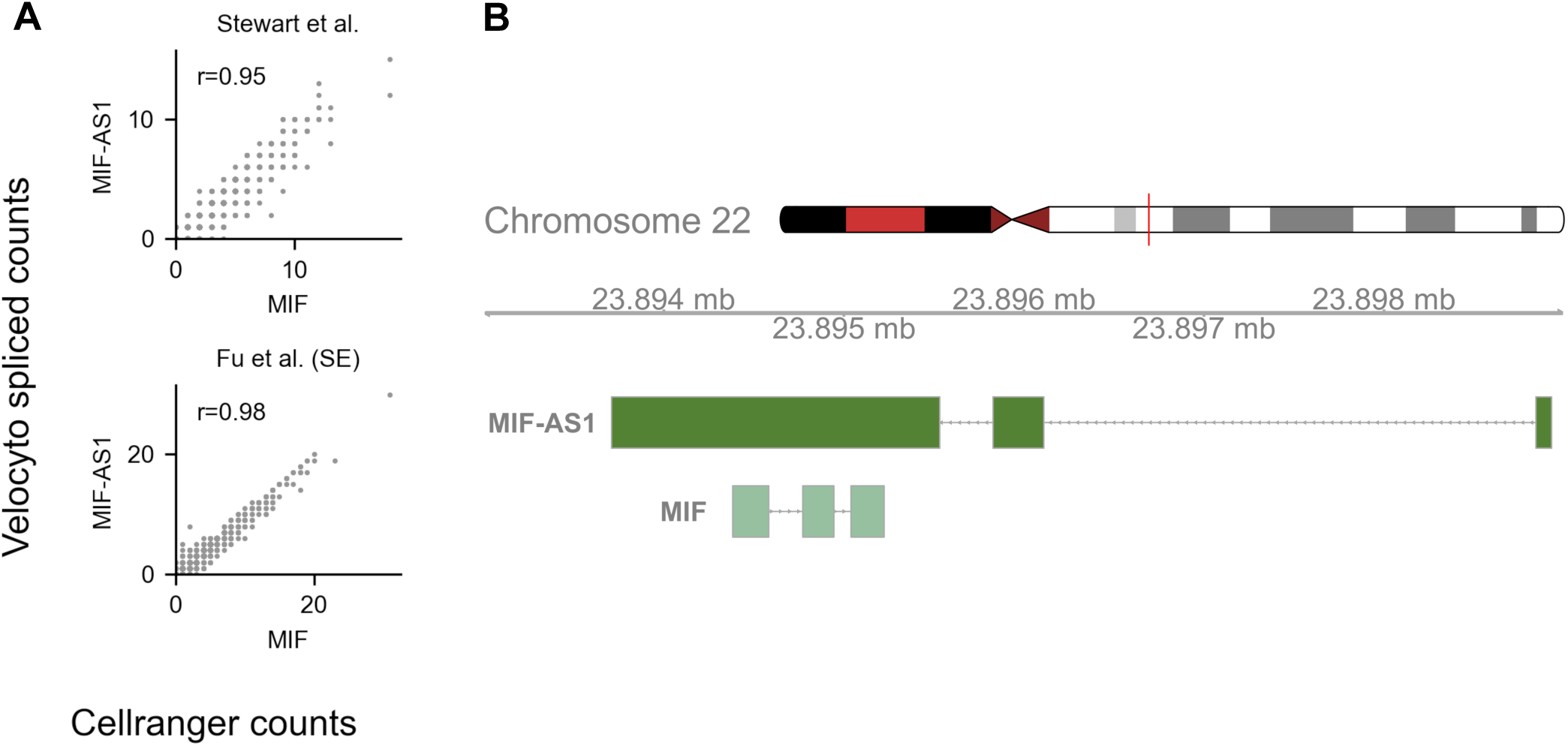
Incorrect assignment of spliced counts by velocyto. (A) Cell Ranger counts of *MIF* vs. velocyto unspliced counts of *MIF-AS1* in the datasets from Stewart *et al*. (upper panel) and Fu *et al*. (lower panel) with Pearson correlation (r). (B) Visualisation of overlap between *MIF* (forward strand) with exon of *MIF-AS1* (reverse strand) on human Chromosome 22.

**Fig. S6.**
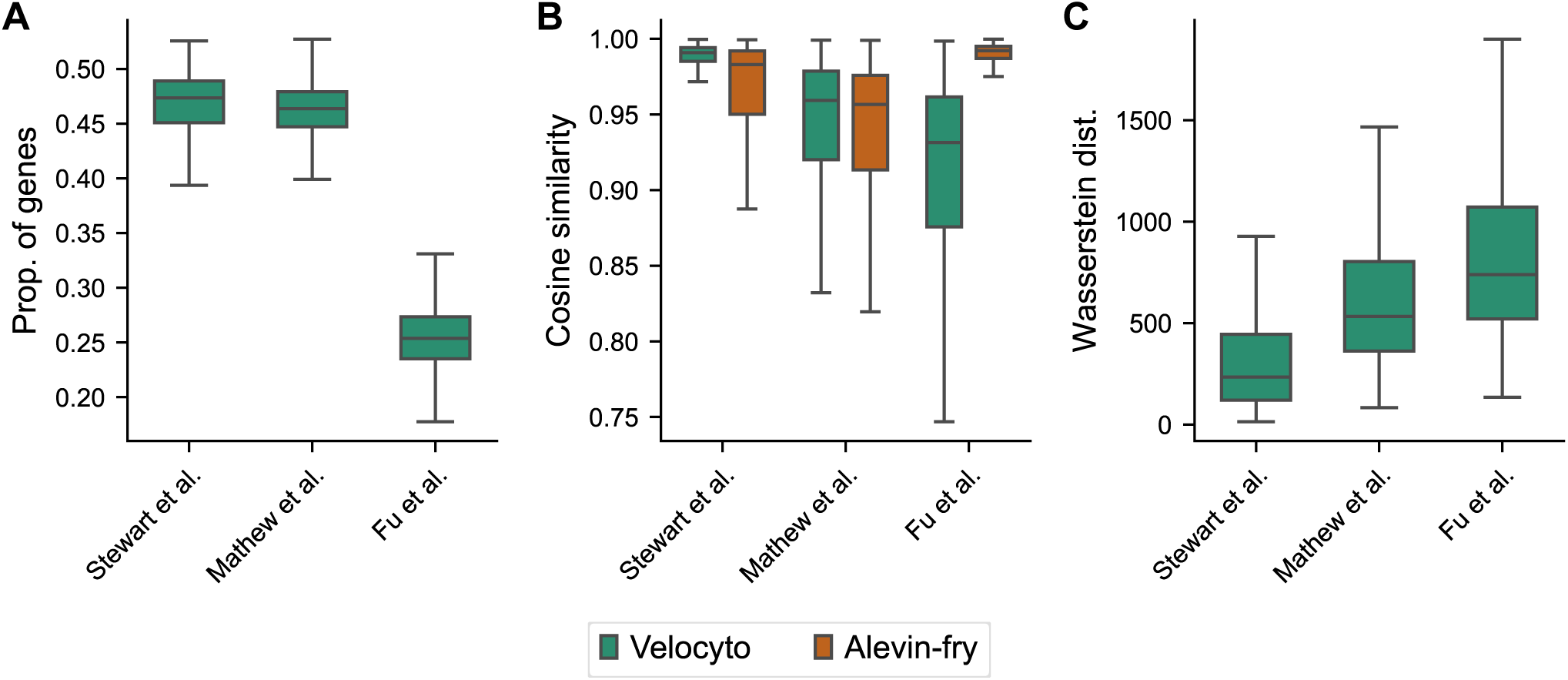
Significance of low similarity between velocity vectors from velocyto and alevin-fry. (A) Proportion of genes per velocity vector with opposing velocity signs between velocyto and alevin-fry. (B) Cosine similarity of velocity vector of each cell with velocity vector of nearest neighbor cell. Nearest neighbor was defined as cell with smallest distance in the respective neighboring graph. (C) Wasserstein distance of transition probability distributions per cell, calculated based on counts from velocyto vs. alevin-fry. Here, transition probability distributions were estimated by using identical velocity genes. Velocity genes were obtained manually as the union of genes selected by velocyto and alevin-fry.

**Fig. S7.**
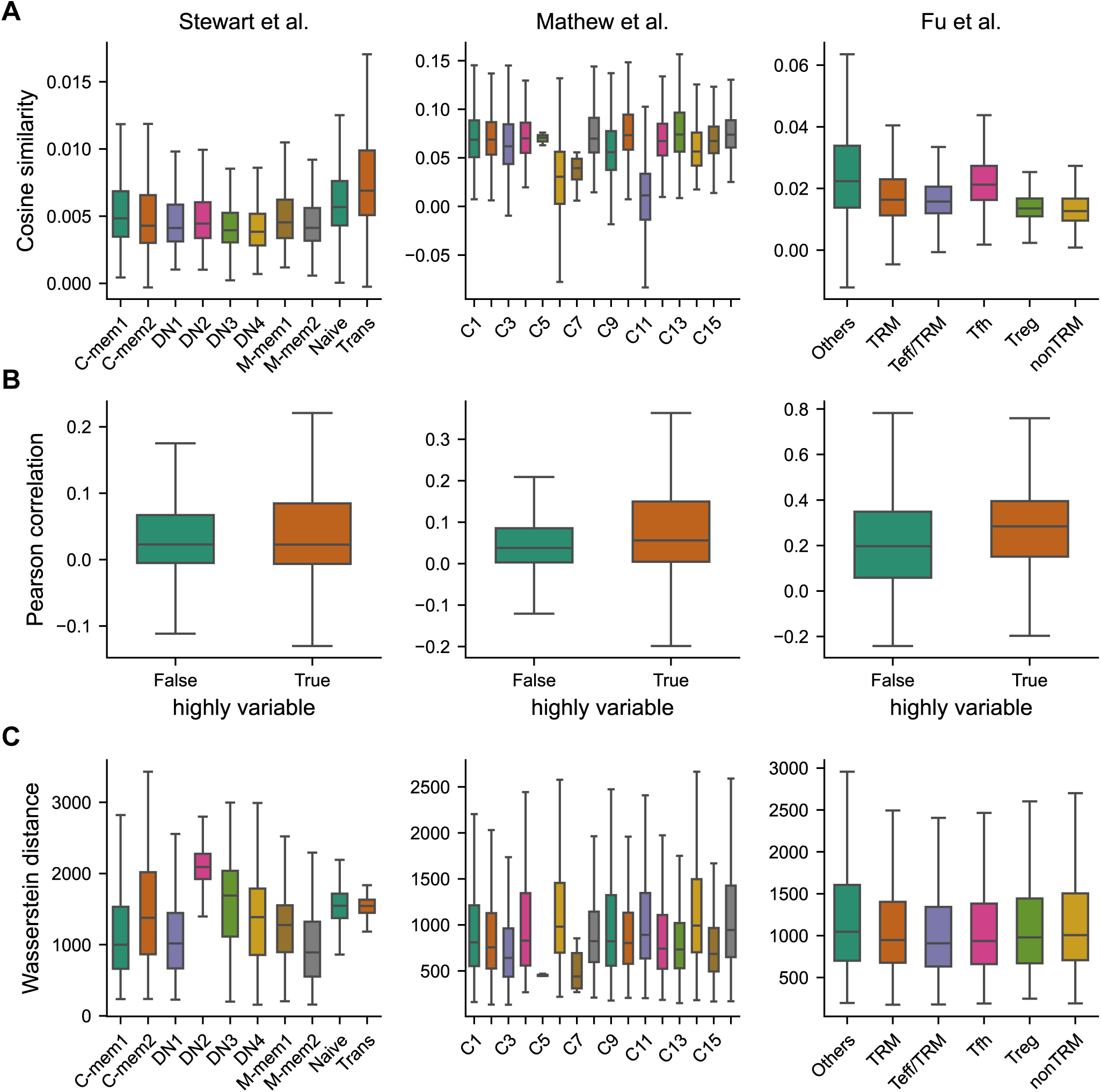
Similarity metrics of velocities are consistent across clusters. (A) Cosine similarity of velocity vectors per cell, divided by cluster. (B) Pearson correlation of velocities per gene, divided by high variability. (C) Wasserstein distance of transition probability distributions per cell, divided by cluster.

**Fig. S8.**
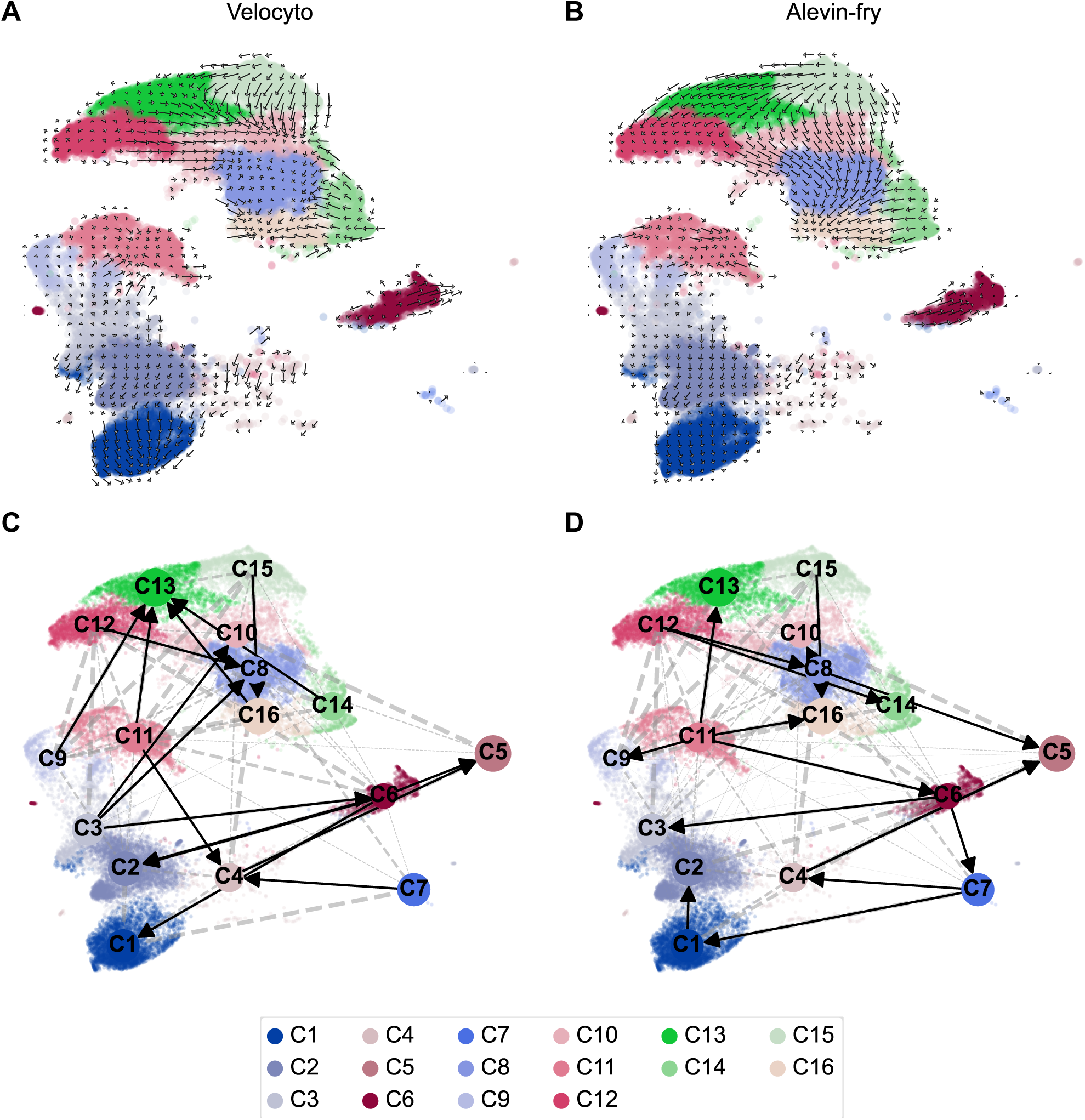
Mathew *et al*.: Inconsistent 2-dimensional projections of velocities. (A,B) Visualisation of velocities from Mathew *et al*. data as gridlines in UMAP-embedding, with counts obtained from velocyto (A) vs. alevin-fry (B). (C,D) PAGA cluster relations projected on UMAP-embedding, with counts obtained from velocyto (C) vs. alevin-fry (D).

**Fig. S9.**
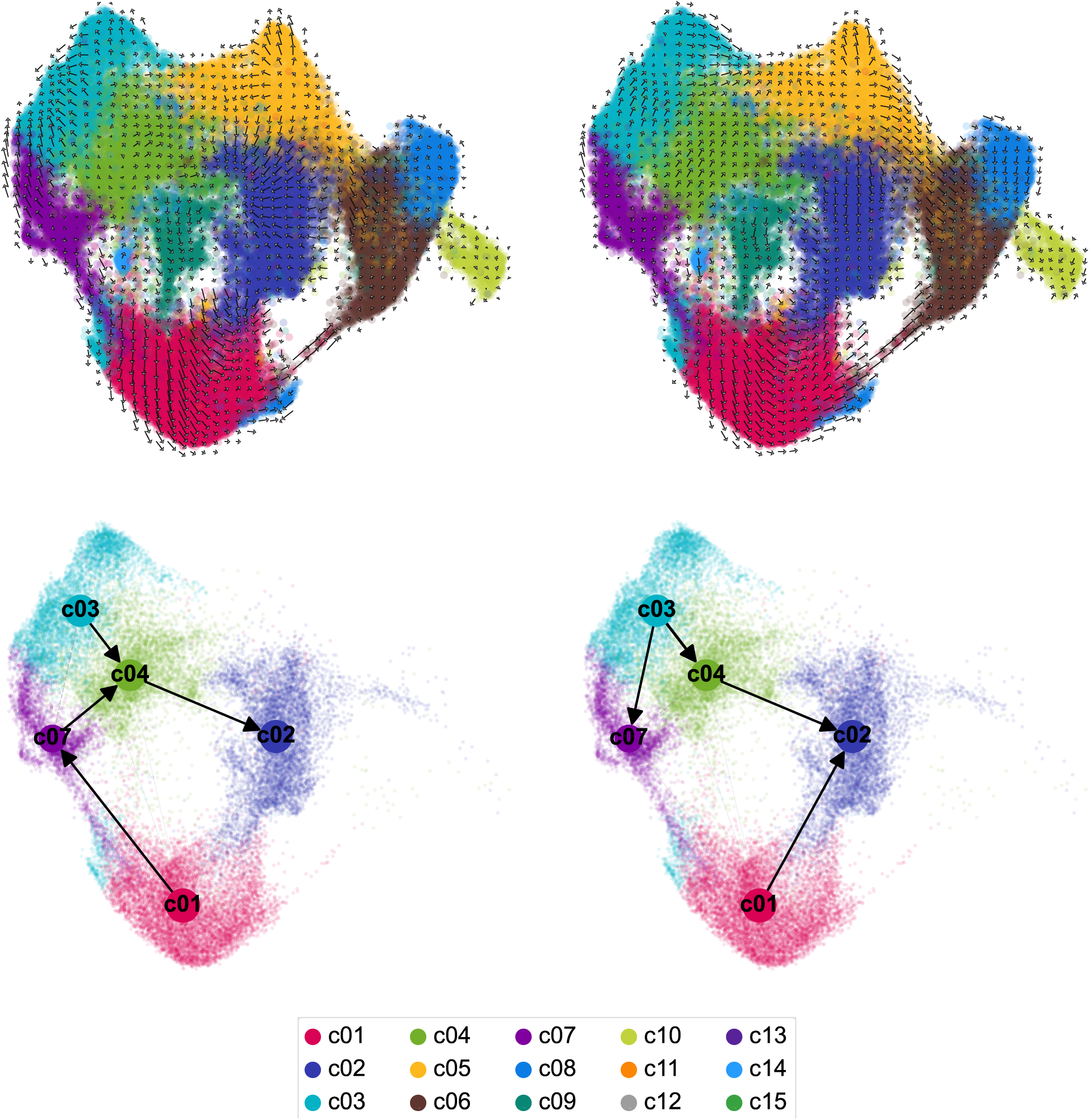
Fu *et al*.: Inconsistent 2-dimensional projections of velocities. (A,B) Visualisation of velocities from Fu *et al*. data as gridlines in UMAP-embedding, with counts obtained from velocyto (A) vs. alevin-fry (B). (C,D) PAGA cluster relations projected on UMAP-embedding, with counts obtained from velocyto (C) vs. alevin-fry (D), showing selected groups (c01, c02, c03, c04 and c07), as described by Fu et al. [2023].

## Notes

### Competing Interest Statement

The authors have declared no competing interest.

